# Tissue transcriptomics of endomyocardial biopsies reveals widespread molecular perturbations independent of leukocyte-rich foci in human myocarditis

**DOI:** 10.1101/2025.07.11.664335

**Authors:** Charles D. Cohen, Jiali H. He, Kevin C. Bermea, Sylvie T. Rousseau, Adam Luo, Obialunanma V. Ebenebe, Kelly Casella, Emanuele Palescandolo, Kory J. Lavine, Eugene Shenderov, Marc K. Halushka, Luigi Adamo

## Abstract

**Background:** Myocarditis is an inflammatory disease of the myocardium, classically defined and graded by histologic criteria that emphasize immune infiltrates and focal cardiomyocyte injury. The broader transcriptional landscape and intercellular signaling networks underlying human myocarditis, particularly among non-immune cells, remain poorly understood.

**Methods:** We performed integrated spatial transcriptomic profiling of 38 endomyocardial biopsy (EMBx) specimens using two complementary platforms: 10X Visium FFPE and GeoMx Digital Spatial Profiling (DSP). The cohort included cases of histologically confirmed myocarditis, borderline myocarditis, and controls. For 10X Visium, data was refined by excluding leukocyte-enriched spots and enriching for cardiomyocyte-specific regions based on canonical marker expression. For GeoMx, immunohistochemistry-guided segmentation enabled targeted transcriptomic analysis of disparate cardiac cellular compartments. Differential gene expression was analyzed independently for each platform and subsequently integrated. These results were further leveraged to infer molecular interaction networks and ligand–receptor relationships in myocarditis relative to controls.

**Results:** Both platforms revealed widespread gene expression changes consistent with immune activation in myocarditis and borderline myocarditis, particularly within cardiomyocyte-enriched regions. These included upregulation of *HLA-A*, *HLA-DQA1*, *B2M*, and *CD74* in myocarditis, consistent with activation of major histocompatibility complex (MHC) class I and II related pathways. Molecular interaction analysis identified *STAT1* and *ISG15* as likely central immune signaling nodes. Ligand–receptor inference highlighted *HLA-A*, *HLA-E*, and *HLA-DQA1* as key receptor hubs interacting with immune ligands such as *IFNG*, *CD8A*, and several members of the (NK) killer-cell immunoglobulin-like receptor (KIR) family.

**Conclusions:** Our findings demonstrate that human myocarditis is characterized by widespread transcriptional dysregulation beyond immune cell foci, including upregulation of genes typically associated with professional antigen-presenting cells in cardiomyocytes. These insights extend our current understanding of myocarditis pathophysiology and suggest new opportunities for its diagnosis and therapeutic targeting.

## INTRODUCTION

Myocarditis is an inflammatory disease of the cardiac muscle that results in over 300,000 deaths annually, which can arise from diverse etiologies, including viral infections, autoimmune reactions, and toxins^1,2^. Among these, viral infections are believed to be the most common cause in developed countries^3^. Clinical presentation of myocarditis is highly variable, ranging from subclinical cases to fulminant heart failure (HF)^4^.

Historically, myocarditis has been considered a spatially heterogeneous disease of the myocardium, histologically diagnosed by discrete foci of immune infiltrates and associated cardiomyocyte injury^4–6^. The presence of immune infiltrates without detectable myocyte damage was classified as ‘borderline myocarditis’ by the Dallas criteria, while the term myocarditis is reserved for cases where immune infiltrates coincide with cardiomyocyte injury^7,8^. However, the broader transcriptional landscape and intercellular signaling networks underlying human myocarditis—and the biological distinctions between myocarditis and borderline myocarditis—remain poorly characterized. Diagnosis of borderline myocarditis remains particularly fraught, as it may reflect either a peripheral glimpse of true myocarditis or a misclassification of a benign immune cell infiltrate.

Emerging insights from cardiac immunology suggest that the heart is intricately connected to the immune system, and that non-immune cells actively participate in immune responses^9^. Therefore, we hypothesized that myocarditis, rather than being a ‘patchy’ disease confined to sites of immune infiltration, represents a global dysfunction of the myocardium, with shared features across etiologies^9,10^. To test this hypothesis, we applied spatial transcriptomic profiling, enabling for interrogation of molecular signaling across regions with and without leukocyte infiltration.

We analyzed endomyocardial biopsies from patients with myocarditis, borderline myocarditis, and appropriate controls. Control samples included one biopsy from a patient undergoing evaluation for suspected myocarditis who was found to have a structurally normal heart, as well as biopsies from patients with heart failure with preserved ejection fraction (HFpEF), whereby myocarditis was ruled out of the differential diagnosis.

Our findings reveal that myocarditis is associated with a diffuse molecular signature that extends well beyond areas of immune cell infiltration. Notably, we demonstrate that in human myocarditis, cardiomyocytes exhibit expression of immune-response genes typically considered exclusive to professional antigen presenting leukocytes.

## METHODS

### Sample Collection and Data Acquisition

Endomyocardial biopsies were collected via transjugular approach for the purpose of patient care. Biopsies were collected from patients with heart failure with preserved ejection fraction (HFpEF) to assess their underlying etiology, and from patients with suspected myocarditis to confirm or rule out the diagnosis.

### 10X Genomics Visium FFPE

All biopsies were reviewed by a cardiac pathologist at Johns Hopkins Hospital. Myocarditis and borderline myocarditis were diagnosed based on the Dallas classification system^8^. A total of 38 archived formalin fixed, paraffin embedded (FFPE) samples were analyzed: 13 control (12 HFpEF and 1 non-myocarditis), 15 borderline myocarditis samples, and 10 myocarditis samples. For one myocarditis patient, two EMBx samples were collected at two different time points, initial presentation and two weeks follow-up. Samples were analyzed using two different versions of 10X Genomics Visium Spatial Transcriptomics technology. A total of 8 samples were analyzed prior to the introduction of CytAssist technology, and the remaining samples were analyzed using the CytAssist platform, elaborated below. Spatial transcriptomic data were generated at the Johns Hopkins Single Cell and Transcriptomics Core with one EMBx per capture area. Raw sequencing data files were processed using Space Ranger software (10X Genomics) prior to subsequent analysis. This pipeline aligned the sequencing reads from fastq files to the reference genome *Homo Sapiens* (GRCh38) and quantified the expression of transcripts within each spot. Then, the analyses of processed data were carried out in R studio v4.3.2 using Seurat suite v5.2.1. A total of 16,827 spatial barcoded spots were retained post processing, considering all disease conditions. To visualize transcriptional clustering by disease classification and minimize confounding factors from technical variation, we grouped samples by preparation method. Samples were processed using two spatial transcriptomic workflows: the original Visium protocol (Visium v1; involving direct tissue placement) and the updated protocol incorporating Visium CytAssist technology (Visium v2). Samples analyzed for Visium v1 included 4 myocarditis and 4 control samples. Using Visium CytAssist (v2), 9 control (HFpEF), 6 myocarditis and 15 borderline myocarditis samples were analyzed. Within each technical group, principal component analysis (PCA) was performed, followed by uniform manifold approximation and projection (UMAP) to visualize the unsupervised clustering of sample groups based on gene expression profiles.

### GeoMx Digital Spatial Profiling

GeoMx Digital Spatial Profiling (DSP) was subsequently performed in the Spatial Cancer Research Immunobiology & Therapeutics (SCRIPT) Laboratory and the Johns Hopkins Experimental and Computational Genomics Core to validate transcriptional findings and enable spatially resolved whole-transcriptome gene expression profiling in cardiac tissue. A total of 8 FFPE EMBx samples—comprising 4 myocarditis and 4 HFpEF controls—were analyzed using the GeoMx DSP platform (NanoString Technologies, Human NGS Whole Transcriptome Atlas v1.0). Sections were co-stained with a multiplexed panel of oligonucleotide-conjugated antibodies including markers for nuclei (Syto83), cardiomyocytes (TNNI3), and leukocytes (CD45), with CD45 signal partially overlapping with autofluorescent lipofuscin. Regions of interest (ROIs) were selected based on these markers to differentiate immune-infiltrated and cardiomyocyte-rich areas, guided by immunohistochemical stained tissue architecture. Within each ROI, multiple Areas of Illumination (AOIs) were segmented to capture distinct cellular foci for gene expression profiling. Barcoded RNA probes within each AOI were released via UV photocleavage and sequenced (Illumina NovaSeq 6000) to generate digital gene counts. Raw data were processed using the GeoMx NGS pipeline (v2.3.3.10) and normalized using upper-quartile scaling (Q3). A total of 18,676 genes were detected before quality control (QC). Given control samples lacked sufficient leukocyte infiltration, CD45⁺ AOIs were absent in this group. Therefore, differential gene expression involving leukocyte-rich regions (TNNI3⁻CD45⁺) from myocarditis samples was compared to non-myocyte, non-leukocyte (interstitial) segments (TNNI3⁻CD45⁻) from controls. Segmentation performance was validated by normalized expression of cardiomyocyte marker *TNNI3* and by reference-based deconvolution of cell-type composition across compartments^10^, confirming enrichment of cardiomyocytes in TNNI3⁺CD45⁻ segments (Supplementary Figure 4A). Unsurprisingly, deconvolution of TNNI3⁻CD45⁻ segments revealed substantial cellular heterogeneity, encompassing fibroblasts, endothelial cells, smooth muscle cells, and other stromal populations, highlighting the broad cellular diversity of this compartment (Supplementary Figure 4B). Similarly, deconvolution of TNNI3⁻CD45⁺ segments from myocarditis samples confirmed the expected enrichment of lymphoid and myeloid immune populations (Supplementary Figure 4C), consistent with transcriptionally active leukocyte foci. Reflecting the variability in marker expression and tissue architecture—particularly within the leukocyte-rich and double-negative (TNNI3⁻CD45⁻) compartments—segmentation was inherently constrained by the limitations of mask-based compartment delineation. For GeoMx DSP data, raw sequencing reads were processed using the GeoMx NGS Pipeline v2.3.3.10 (Bruker [NanoString] Technologies). Differential gene expression analyses between experimental groups were performed using linear mixed modeling (LMM) with Benjamini-Hochberg False Discovery Rate (FDR) correction. Cell type proportions across AOIs were estimated via SpatialDecon, using a human immune–cardiac reference profile^11^. All downstream analyses, including data integration with Visium, DEG comparisons, and visualizations (e.g., heatmaps, Venn diagrams), were performed using R v4.3.2 with ggplot2, dplyr, and ComplexHeatmap.

### Data Preprocessing and Quality Control

For 10X Visium, raw transcriptomics data were processed in the standard 10x Genomics Space Ranger pipeline. Filtered feature-barcode matrices were imported into R using the Read10X_h5 function from the Seurat package (v5.2.1), to initiate downstream data processing. QC metrics were evaluated, including the number of unique molecular identifiers (UMIs) per spot and the number of genes detected per spot. For data filtering, barcoded spots with 647 or fewer UMIs (≤ 6.25% [one quarter of the first quartile] threshold) were removed to eliminate low-quality spots or those outside the tissue boundary. This threshold for border filtering was determined from the overall distribution of UMI counts across all samples ranging from 0 to 110,120 (median UMIs = 5,959). To further reduce technical artifacts associated with tissue quality, we generated a curated list of RNA integrity assessment genes (including mitochondrial and RNA degradation markers), for subsequent removal from each sample (Supplementary Table 2). To refine cell-specific gene expression, we employed several filtering strategies to separate technical noise and improve biological signal. First, mitochondrial genes, including MT-ND1, MT-ND2, MT-CO2, and MT-ATP6 (full list in Supplementary Table 2), were removed from all samples to mitigate technical artifacts. After determining broad gene expression between experimental groups (post-QC filtering), spots rich in leukocyte markers were removed using a comprehensive custom curated list (Supplementary Table 2), to focus on non-immune cardiac tissue. The data was then refined by enriching for spots with high counts of cardiomyocyte-specific genes. Spots were considered enriched for cardiomyocytes if they expressed at least 5 of the cardiomyocyte marker genes selected, above a threshold level of expression of 1.5 (Supplementary Table 2). Based on the filtering conditions described above, we generated three analysis-ready datasets through progressive filtering: (1) a QC-filtered dataset excluding mitochondrial genes and suboptimal capture spots, (2) a dataset excluding sequencing spots enriched in immune cell markers, and (3) a cardiomyocyte-enriched dataset of immune-depleted spots.

For GeoMx DSP analysis, raw data was Q3 normalized and grouped into disease conditions (control, myocarditis) and statistically compared using LMM with Benjamini-Hochberg FDR correction.

### Differential Gene Expression Analysis

Differential gene expression was assessed using three independent and complementary approaches with varying rigor: DESeq2 (most stringent), MASTcpmDETrate (most sensitive) and Seurat’s FindAllMarkers (intermediate sensitivity). To identify robust gene expression signatures distinguishing experimental groups (control, myocarditis, and borderline myocarditis) we evaluated concordance across the results obtained with these three methods. For each method using 10X Visium data, genes were selected based on significance, effect size, and consistency across comparisons. This analysis was performed across three progressively filtered datasets, as described above: (1) a QC-filtered dataset, (2) an immune cell–depleted dataset, and (3) a cardiomyocyte-enriched subset derived from immune-depleted data. This stepwise approach enabled the identification of both shared and cell type–enriched transcriptional signatures across cardiac tissue compartments. Further details are provided below. Fold change (FC) thresholds were empirically determined for each dataset based on the statistical distribution, variability inherent to each dataset, and the sensitivity of each differential expression method. Given that DESeq2, MASTcpmDETrate, and Seurat’s FindAllMarkers differ in stringency and data handling, the dynamic range and number of differentially expressed genes (DEGs) varied across methods. To ensure meaningful biological interpretation while minimizing noise, we selected FC thresholds that balanced gene list size with statistical confidence.

#### DESeq2

For DESeq2 analysis (v1.42.1), raw count matrices were first filtered to remove genes detected in <2 spots, ensuring sufficient expression for reliable statistical modeling. Subsequently, to address zero values/counts (due to sparsity from DESeq2 filtering), a pseudocount of 1 was added to all gene counts to facilitate accurate log-transformation and variance stabilization^12^. For each comparison, the relevant disease group was set as the primary variable in the design formula to guide differential gene expression analysis. The group designated as the variable of interest (as well as the denominator) varied depending on the comparison being made. Genes were classified as differentially expressed if they met both an adjusted p-value <0.001 (Benjamini-Hochberg False Discovery Rate [FDR] correction) and an absolute log_2_ FC threshold >1.5.

#### MASTcpmDETrate

Raw count data were normalized to counts per million (CPM) and log_2_-transformed. Next, we tailored a zero-inflated model with disease condition as predictor variable, as implemented in the MAST framework (v.1.28.0)^13^. Genes that had non-zero expression in >10% of spots in at least one of the experimental groups were considered in this analysis. This test is a modification of ‘MAST’ considering cellular detection rate as a covariate, which has been successfully performed in a benchmark study for detecting differential expression from scRNA-seq^14^. Genes with an adjusted p-value <0.001 (Benjamini–Hochberg FDR correction) were considered statistically significant. Given this method is the most sensitive approach in our pipeline, adaptative log_2_ FC thresholds were set for each of the analyzed datasets. The log_2_ FC threshold was computed for each dataset to maximize the trade-off between statistical stringency and the resulting gene set size. For the QC-filtered dataset log_2_ FC threshold of >3 was applied. For the subsequent immune-independent dataset, we set an absolute log_2_ FC threshold of >2.5, to retain sufficient DEGs given the moderate sample size and mixed cell population. For the further refined cardiomyocyte-enriched dataset derived from immune-depleted spots, we applied the same threshold (>2.5 absolute log_2_ FC).

#### FindAllMarkers (Seurat)

Log-transformed, normalized data were analyzed using the Wilcoxon rank-sum test with a minimum detection threshold of 10% of spots and a baseline log_2_ FC threshold cutoff of 0.25, as typically used in this pipeline^15^. The results were then further filtered by adjusted p-value <0.001 and adaptive absolute log_2_ FC cut-offs. The log_2_ FC threshold was computed for each dataset to maximize the trade-off between statistical stringency and the resulting gene set size.

To account for differences in signal-to-noise ratio across datasets, we applied empirically guided FC thresholds. For the QC-filtered dataset, an absolute log_2_ FC threshold of >2 was used. For the subsequent immune-independent dataset, a threshold of absolute log_2_ FC > 2 was required. For the further refined cardiomyocyte-enriched dataset of immune-depleted spots, an absolute log_2_ FC >1 threshold was applied.

For GeoMx DSP analysis, differential gene expression between experimental groups was performed using LMM with Benjamini-Hochberg FDR correction, as discussed above. A total of 11,430 transcriptomic targets were captured post QC for LMM differential expression analysis within relevant experimental groups.

### Protein-protein interactions and ligand**–**receptor inference

To investigate immune-related signaling pathways and interactions relevant to myocarditis, we utilized two curated databases of genes, proteins, and molecular networks. InnateDB^16^ was used to infer intracellular protein–protein interactions (PPI) involved in the innate immune response, and OmniPath^17^ was employed to compute ligand–receptor (L-R) networks to visualize intra- and intercellular signaling.

#### InnateDB Analysis

This analysis enabled the identification of immune mechanisms that are likely to distinguish myocarditis from controls, highlighting key pathways and mediators associated with inflammatory activation in cardiac tissue. The top 150 DEGs from FindAllMarkers from Visium data with leukocytes removed and cardiomyocytes enriched were selected and cross-referenced with curated interactions from InnateDB to interrogate the PPI. The InnateDB dataset was filtered to retain only human proteins, and processed to remove duplicate and non-standard gene identifiers, using dplyr (v1.1.4) and tidyr (v1.3.1). Only interactions involving at least one gene from the input list were retained. The final PPI network was visualized using the igraph package (v2.1.4), with node color assigned via RColorBrewer (v1.1.3) to reflect topological heterogeneity and node size scaled by degree of connectivity (Supplementary Figure 2). Interactions were visualized as undirected edges, suggesting the interconnected role of immune signaling proteins in myocarditis.

#### OmniPath Analysis

DEGs from the QC-filtered dataset were further analyzed using the OmniPath database—a literature-curated resource of signaling pathways and molecular interaction networks. Ligand–receptor interactions were retrieved through the OmniPath database, using the import_intercell_network function from OmniPathR package (v3.10.1). Directional interactions were filtered to retain only those with a ligand as the source and a receptor as the target. Subsequently, a directed graph was constructed to visualize interactions using chord diagrams in the circlize package (version 0.4.16). For the HLA-focused analysis (Figure 5), interactions were filtered to retain only those involving overlapping DEGs between GeoMx and Visium platforms. A custom set of 21 HLA genes present in OmniPath—including HLA-A, HLA-E, and HLA-DQA1—was manually defined and highlighted. All other interactors were displayed in grey to emphasize HLA-centric signaling. Chord width reflects the number of unique ligand–receptor interactions involving each gene.

### Ingenuity Pathway Analysis (IPA)

DEGs identified between myocarditis and control were subjected to IPA (QIAGEN) to assess pathway enrichment. IPA was performed on the DEGs derived from FindAllMarkers throughout the study, as this approach balanced sensitivity and stringency, capturing robust yet biologically interpretable DEG sets without over-inflating gene lists. The DEG list used for IPA was filtered for significance based on an adjusted p-value <0.05 and absolute log_2_ FC >1.5. IPA core analysis was performed with default settings, restricting the analysis to human genes and experimentally observed relationships. Canonical pathways were prioritized based on z-score (reflecting pathway activation or inhibition) and p-value (Fisher’s exact test, displayed as -log_10_ p-value). Pathways with an absolute z-score ≥1.3 and p <0.05 were considered significant.

## RESULTS

### Clinical characteristics of study participants and overall study design

We performed tissue transcriptomics analysis of 38 EMBx samples collected for clinical care purposes (Figure 1A). Of these, 10 samples had a pathological diagnosis of myocarditis (including immune checkpoint inhibitor (ICI)–associated myocarditis [n=2], eosinophilic myocarditis [n=1], and lymphocytic myocarditis [n=7]). Fifteen samples were diagnosed as borderline myocarditis, while 13 samples from control patients were collected either to rule out myocarditis [n=1] or to gain insights into the etiology of heart failure with preserved ejection fraction (HFpEF, [n=12]). Myocarditis patients exhibited reduced left ventricular ejection fraction (LVEF) and elevated NT-proBNP levels (Supplementary Table 1). Median LVEF was 37.5% in myocarditis, 65% in controls, and 58.8% in borderline myocarditis. Similarly, median NT-proBNP levels were 7,931 pg/mL in myocarditis, 337 pg/mL in controls, 4,535 pg/mL in borderline myocarditis. Complete clinical and diagnostic characteristics are provided in Supplementary Table 1.

**Figure 1:**
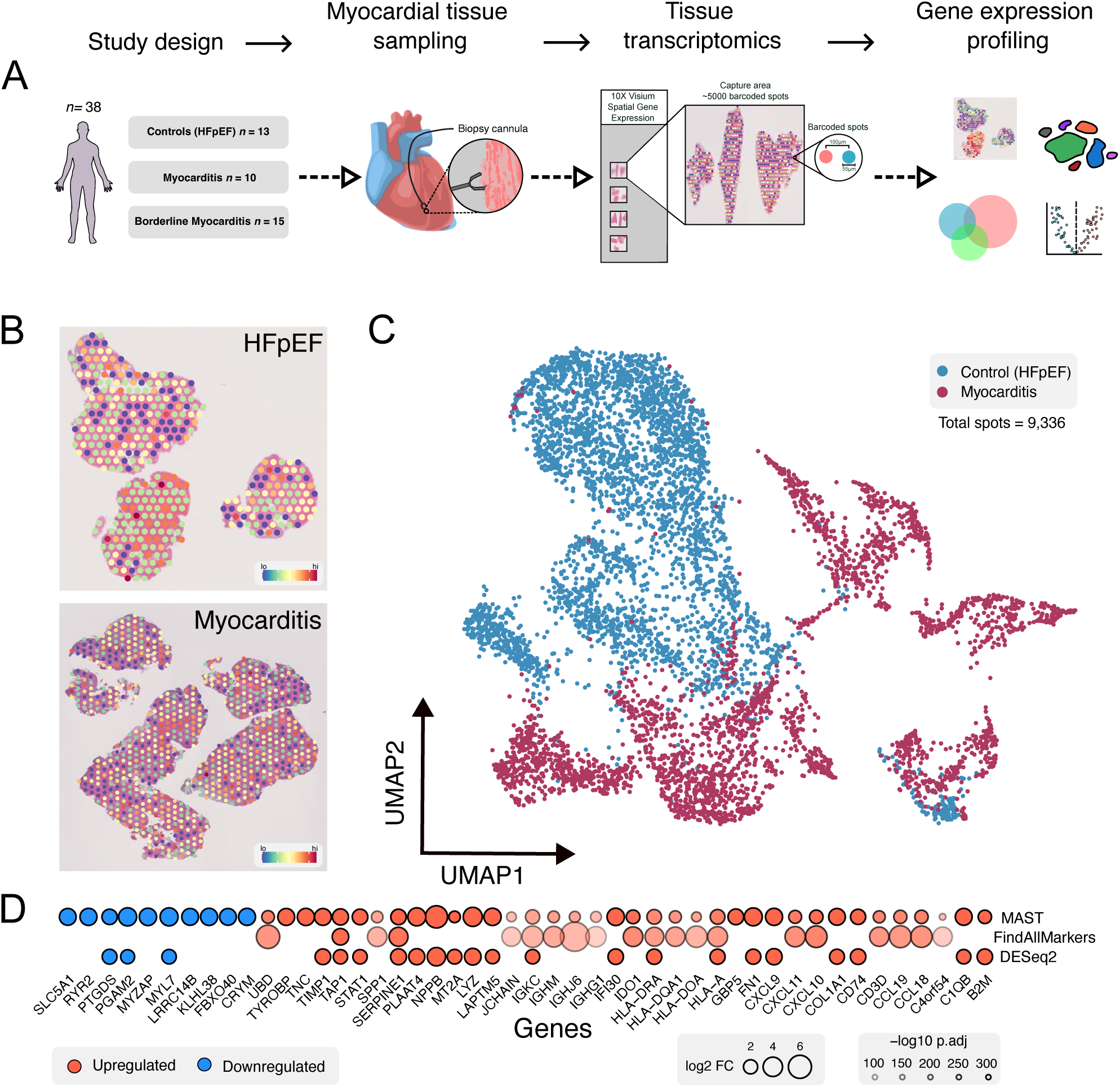
Spatial transcriptomic profiling of EMBx reveals distinct gene expression patterns in clinical myocarditis compared to controls. **(A)** Schematic of study design: EMBx were collected from patients with myocarditis (*n* = 10), controls (HFpEF; *n* = 13), and borderline myocarditis (*n* = 15), followed by *in silico* analysis and independent validation. **(B)** Representative spatial feature plots showing heterogeneous gene expression in myocarditis and HFpEF controls. **(C)** UMAP projection of all spatial barcoded spots (spots = 9,336) demonstrating clear separation between HFpEF controls (blue) and myocarditis (maroon). **(D)** Differentially expressed genes (DEGs) identified using three independent methods with increasing stringency (MASTcpmDETrate, Seurat’s FindAllMarkers, DESeq2). Circle size reflects log_2_ fold change (FC), circle shading (alpha) corresponds to statistical significance (–log₁₀ adjusted p-value [p.adj]); circle color indicates direction of regulation (red = upregulated, blue = downregulated, denominator = control).

### Tissue transcriptomic profiling of myocarditis and control samples indicates widespread and distinct gene expression patterns

To explore the transcriptional characteristics of myocarditis, we first focused on data collected from myocarditis samples and controls. 10X Visium-based analyses of these samples produced 9,336 barcoded spatial transcriptomics ‘spots’ following QC filtering (Figure 1A&B). UMAP plots highlighted distinct clustering of the sequencing spots from these two patient groups, suggesting that myocarditis is characterized by widespread dysregulation of gene expression (Figure 1C). Subsequently, differential gene expression between myocarditis and control sequencing spots was performed using three strategies—MASTcpmDETrate, DESeq2, and FindAllMarkers—to determine the top common DEGs consistently identified across strategies (Figure 1D, Supplementary Table 3). Examination of these top dysregulated genes identified genes previously implicated in myocarditis, as well as novel genes not previously described in myocarditis (Figure 1D, Supplementary Table 3). Known genes associated with myocarditis included *CD3D*^18^, *CXCL9*^19^ and *COL1A1*^20^, all of which were significantly augmented in myocarditis relative to control samples (Figure 1D, Supplementary Table 3). Unexpectedly, several genes related to antigen presentation were significantly upregulated. These included *HLA-A* and *HLA-DRA*; critical components of human MHC class I and MHC class II complexes, respectively. Collectively, these findings highlighted a robust and widespread immune activation signature in myocarditis, characterized by enhanced expression of antigen presentation genes.

### Refining tissue transcriptomics data by excluding leukocyte-rich spots while including cardiomyocyte-enriched foci

Next, we refined the data by removing all sequencing spots exhibiting leukocyte-rich gene signatures, to explore whether we could detect a generalized gene expression signature of myocarditis that was not driven by immune infiltrates. Removal of leukocyte-rich areas yielded 3,966 spots, which showed distinct separation between myocarditis and controls via UMAP (Figure 2A). Unsurprisingly, collagen genes were upregulated in myocarditis after data refinement, including *COL1A1* and *COL3A1*^20^. Moreover, genes encoding cardiac structural proteins, including *MYH7* and *RYR2* were significantly downregulated (Figure 2B, Supplementary Table 4). Interestingly, several immunoglobulin and HLA-related genes remained upregulated in myocarditis, relative to controls (Figure 2B, Supplementary Table 4). Among these included *IGKC*, *HLA-A*, *IGHJ6* and *IGHG1*, which remained significantly elevated in myocarditis, despite the exclusion of leukocyte-rich spots. Next, we performed IPA on DEGs identified from FindAllMarkers (Figure 2C, Supplementary Table 5). IPA indicated enrichment of gene programs related to upregulation of *Interferon gamma signaling* and *MHC class II antigen presentation*, underscoring that upregulation of components of the inflammatory response in myocarditis persist despite crude removal of leukocyte signatures.

**Figure 2:**
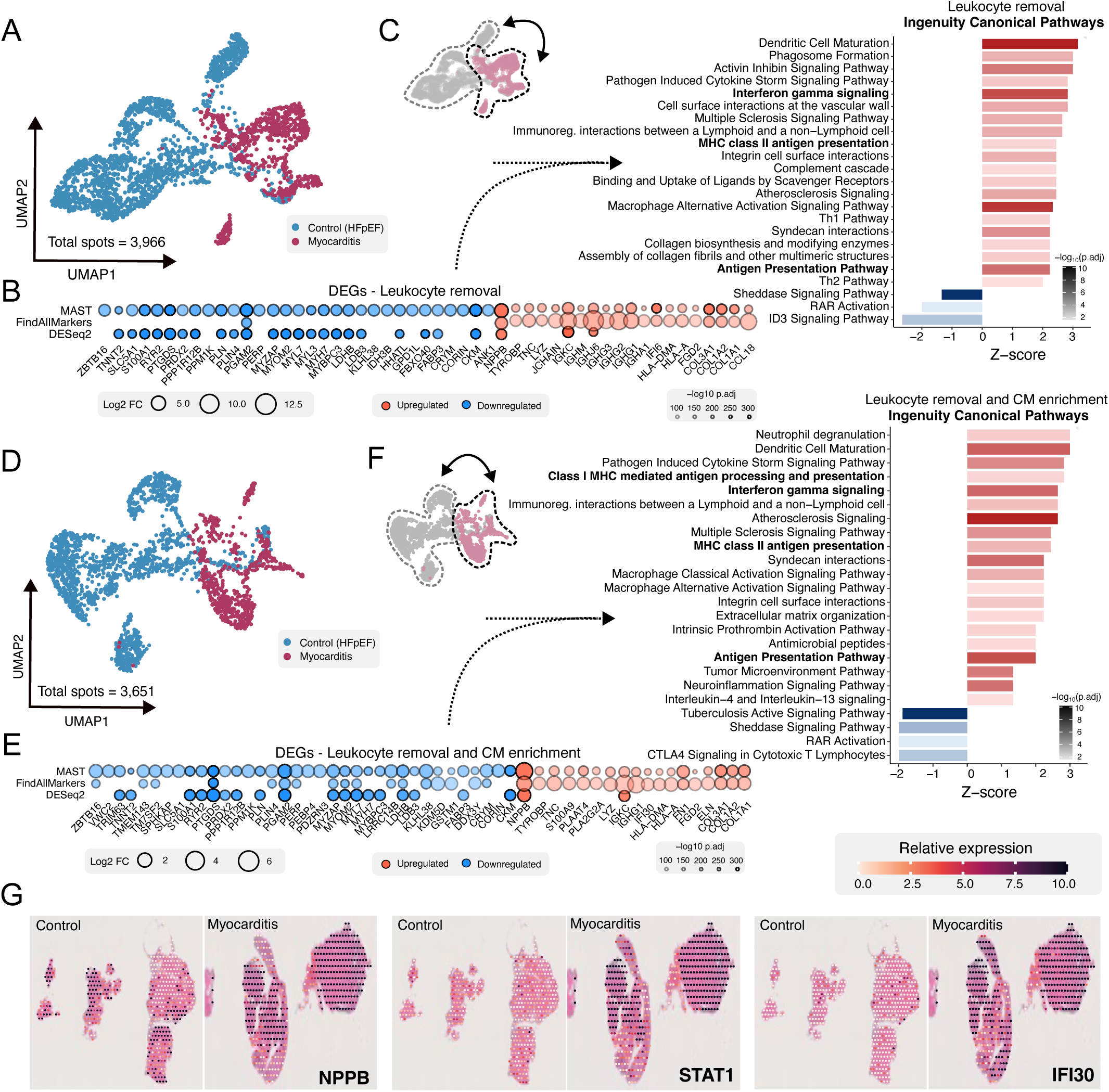
Refining tissue transcriptomics by omitting immune-rich sequencing spots and isolating cardiomyocyte-enriched regions. **(A)** Representative UMAP illustrating distinct clustering between control and myocarditis patients, after removal of immune-rich sequencing spots. **(B)** Top 50 DEGs detected between three DEG strategies. **(C)** IPA performed on DEGs identified by FindAllMarkers. **(D)** Representative UMAP showing distinct clustering of HFpEF controls and myocarditis patients, post removal of immune-rich spots and subsequent enrichment of cardiomyocyte genes. **(E)** Top 50 DEGs using each DE strategy, with immune foci removed and CMs enriched. Circle size reflects log_2_ fold change (FC), circle shading (alpha) corresponds to statistical significance (–log₁₀ adjusted p-value [p.adj]); circle color indicates direction of regulation (red = upregulated, blue = downregulated, denominator = control). (**F**) IPA performed on DEGs post removal of immune-rich spots and subsequent enrichment of cardiomyocyte genes. **(G)** Spatial transcriptomic visualization of selected differentially expressed genes in myocarditis versus control hearts. Representative tissue sections show relative expression of *NPPB*, *STAT1*, and *IFI30*, overlaid on H&E-stained images. Expression intensity is scaled per gene and color-coded according to the accompanying heat scale (orange = low, black = high).

To further distil the data, we enriched sequencing spots for cardiomyocyte-specific genes after excluding leukocyte-rich spots. Here, we identified a total of 3,651 sequencing spots, which still showed distinct clustering between myocarditis and controls (Figure 2D). Differential gene expression analysis was performed synonymously, highlighting similarly expressed collagen genes, cardiac structural genes and *NPPB*—which was increased by a greater magnitude than after leukocyte removal alone (Figure 2E, Supplementary Table 6). Despite leukocyte removal and cardiomyocyte enrichment, several immune-related genes—including HLA and immunoglobulin transcripts—remained upregulated (Figure 2E, Supplementary Table 6), aligning with IPA results showing enrichment of *Interferon gamma signaling*, *Class I MHC antigen presentation* and *MHC class II antigen presentation* (Figure 2E-F, Supplementary Table 7). In addition, Protein-Protein Interaction (PPI) networks were computed via InnateDB^16^ on leukocyte-depleted, cardiomyocyte-enriched DEGs, which emphasized that *STAT1* and *ISG15* are likely central molecular hubs that regulate these processes in myocarditis (Supplementary Figure 2). Together, this reinforces the observation that components of the antigen presentation pathway are transcriptionally active in non-immune cardiac cells in the context of myocarditis. Pertinent genes were subsequently spatially mapped to initial EMBx micrographs, which demonstrated widespread upregulation of genes including *NPPB*, *STAT1* and *IFI30* in myocarditis relative to controls (Figure 2G).

### Incorporation of borderline myocarditis samples reveals transcriptional heterogeneity across the diagnostic spectrum

To assess whether borderline myocarditis represents a transitional or distinct molecular state from myocarditis, we analyzed EMBx from patients classified as ‘borderline’, based on the Dallas criteria. First, we considered all samples by illustrating all existing spots by UMAP (total spots = 16,827). This visualization displayed scattered, diffuse clustering between control, borderline myocarditis and myocarditis (*n* = 15, Figure 3A). Most borderline myocarditis samples displayed an intermediate or distinct transcriptional landscape via UMAP (*n* = 10, Figure 3B). A subset of borderline myocarditis samples showed notable transcriptomic alignment with controls or myocarditis (Figure 3C). These borderline myocarditis samples were removed from further analysis due to potential histopathological misclassification (*n* = 5, Supplementary Figure 3). Next, transcriptomic spots were mapped onto a revised UMAP, which showed more distinct clustering of borderline myocarditis samples (Figure 3D). Borderline myocarditis samples were then compared via differential expression relative to controls (Figure 3E, Supplementary Table 8) and myocarditis (Figure 3F, Supplementary Table 8) to assess the molecular diversity of borderline myocarditis. Borderline myocarditis showed predominantly upregulated, but also several downregulated genes compared to controls (Figure 3E, Supplementary Table 8). Given that controls in this study were almost exclusively HFpEF, the observed upregulation of *MYH7*, *NPPA* and *COL1A2* was consistent with the expected transcriptomic profile associated with this phenotype, consistent with both rodent and human data^21–25^ (Figure 3D, Supplementary Table 8). Several novel genes were also upregulated in controls relative to borderline myocarditis, including *ITM2B*, an amyloid-associated gene^26^; *GPD1L*, involved in oxidative metabolism and cardiac arrythmias^27^; and *PSAP*, a regulator of lysosomal and sphingolipid metabolism^28^ (Figure 3E, Supplementary Table 8), which suggests that HFpEF controls resemble a more structurally and metabolically perturbed state, compared to borderline myocarditis. In contrast, myocarditis samples display upregulated inflammatory genes, such as *CD74*, *B2M* and *C1QC* (Figure 3F, Supplementary Table 8), relative to borderline myocarditis—underscoring that the degree of inflammation in diagnosed myocarditis is higher than borderline cases. Moreover, several HLA-encoding genes were upregulated in myocarditis—including *HLA-A*, *HLA-DRA* and *HLA-E* (Figure 3F, Supplementary Table 8)—reinforcing that myocarditis is associated with upregulation of signaling pathways related to antigen presentation, independent of reference group comparison. Removal of leukocyte-rich spots and enrichment for cardiomyocytes confirmed these findings. Indeed, differential gene expression revealed broad upregulation of immune-related genes in myocarditis, including antigen presentation (including *CD74*, *HLA-DPA1,* and *HLA-DRA*) and interferon-related transcripts (including *IFI30* and *GBP1*), as well as inflammatory mediators (including *B2M* and *PSMB8*), further illustrating the widespread transcriptional activation of inflammatory and immunogenic programs in myocarditis compared to borderline myocarditis (Figure 3G, Supplementary Table 8). Taken together, these findings indicate that borderline myocarditis is a heterogeneous diagnosis that includes both samples with an intermediate level of inflammatory activation between controls and myocarditis but without no clear upregulation of antigen presentation related pathways, and samples with a transcriptional signature superimposable to controls or myocarditis.

**Figure 3:**
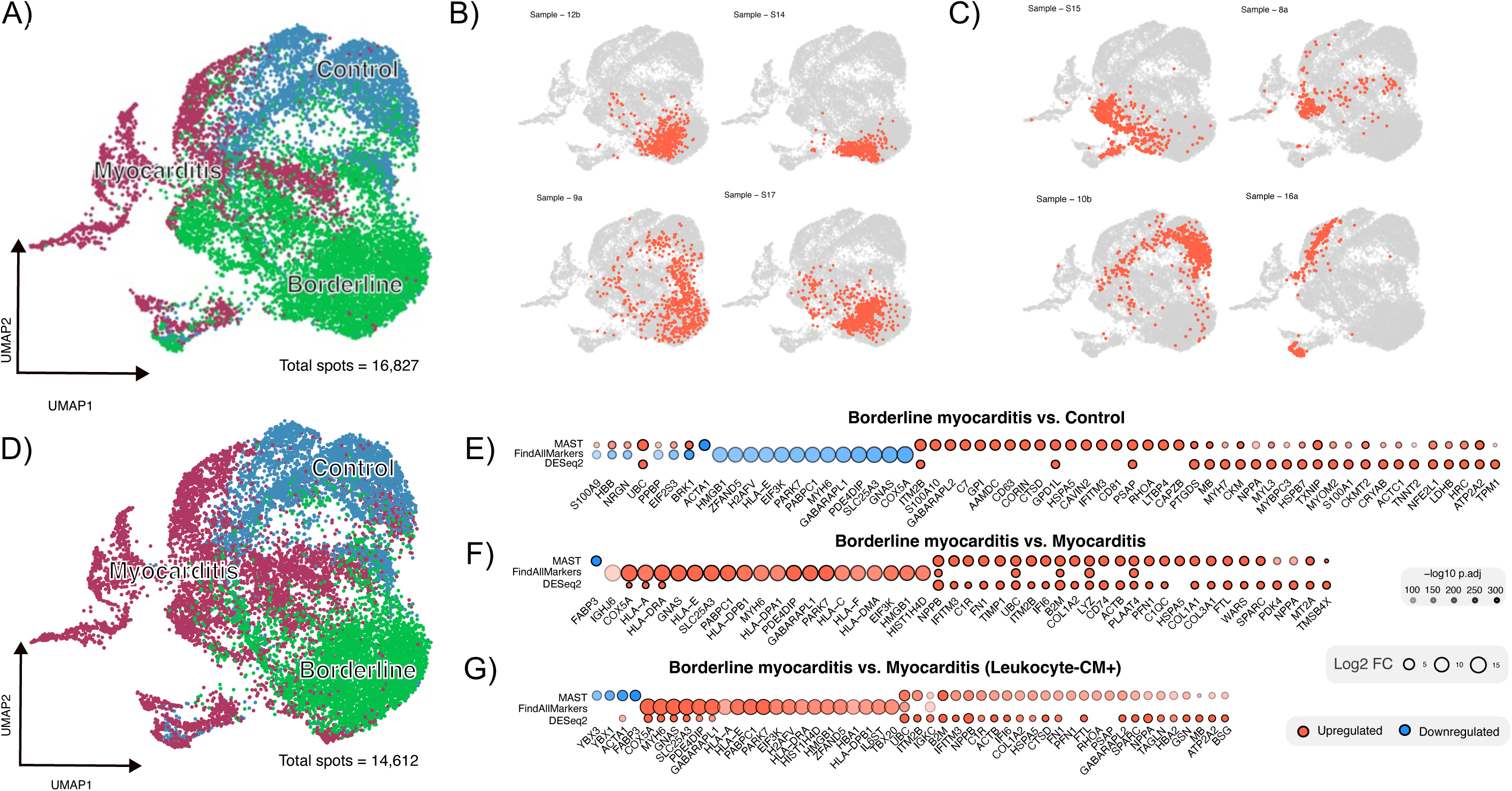
Borderline myocarditis samples exhibit an intermediate transcriptional profile between controls and myocarditis. **(A)** UMAP projection of all captured spots (spots = 16,827), showing transcriptional separation between controls (blue), confirmed myocarditis (maroon), and borderline myocarditis (green). **(B)** UMAP highlighting the spatial distribution of borderline myocarditis included in subsequent analysis. **(C)** UMAP highlighting the spatial distribution of borderline myocarditis misclassified by traditional histological diagnosis. **(D)** Revised UMAP projection of included borderline myocarditis samples (*n* = 10, spots = 14,612), demonstrating more distinct separation of borderline myocarditis samples. **(E)** Differential expression analyses comparing borderline myocarditis (denominator) to controls, **(F)** borderline myocarditis to diagnosed myocarditis, and **(G)** borderline myocarditis to myocarditis within immune-depleted, cardiomyocyte-rich regions. Circle size reflects log2 fold change (FC), circle shading (alpha) corresponds to statistical significance (–log₁₀ adjusted p-value [p.adj]); circle color indicates direction of regulation (red = upregulated, blue = downregulated). Minor discrepancies in directionality reflect methodological differences near significance thresholds.

### Immunohistochemistry-directed spatial transcriptomics corroborates Visium-based analysis

To cross-validate 10X Visium findings, we performed targeted spatial transcriptomics using GeoMx DSP NanoString (Bruker) technology. Cardiac tissue slices from control (*n* = 4) and myocarditis (*n* = 4) samples were stained with three immunohistochemical markers—TNNI3 to delineate cardiomyocytes, CD45 to identify leukocytes, and Syto83 to identify nuclei (Figure 4A). This allowed for segmentation-enabled transcriptomic analysis across three spatially distinct compartments: TNNI3⁺CD45⁻ cardiomyocytes, TNNI3⁻CD45⁺ leukocytes, and TNNI3⁻CD45⁻ non-myocytes (Figure 4B, Supplementary Table 9).

**Figure 4.**
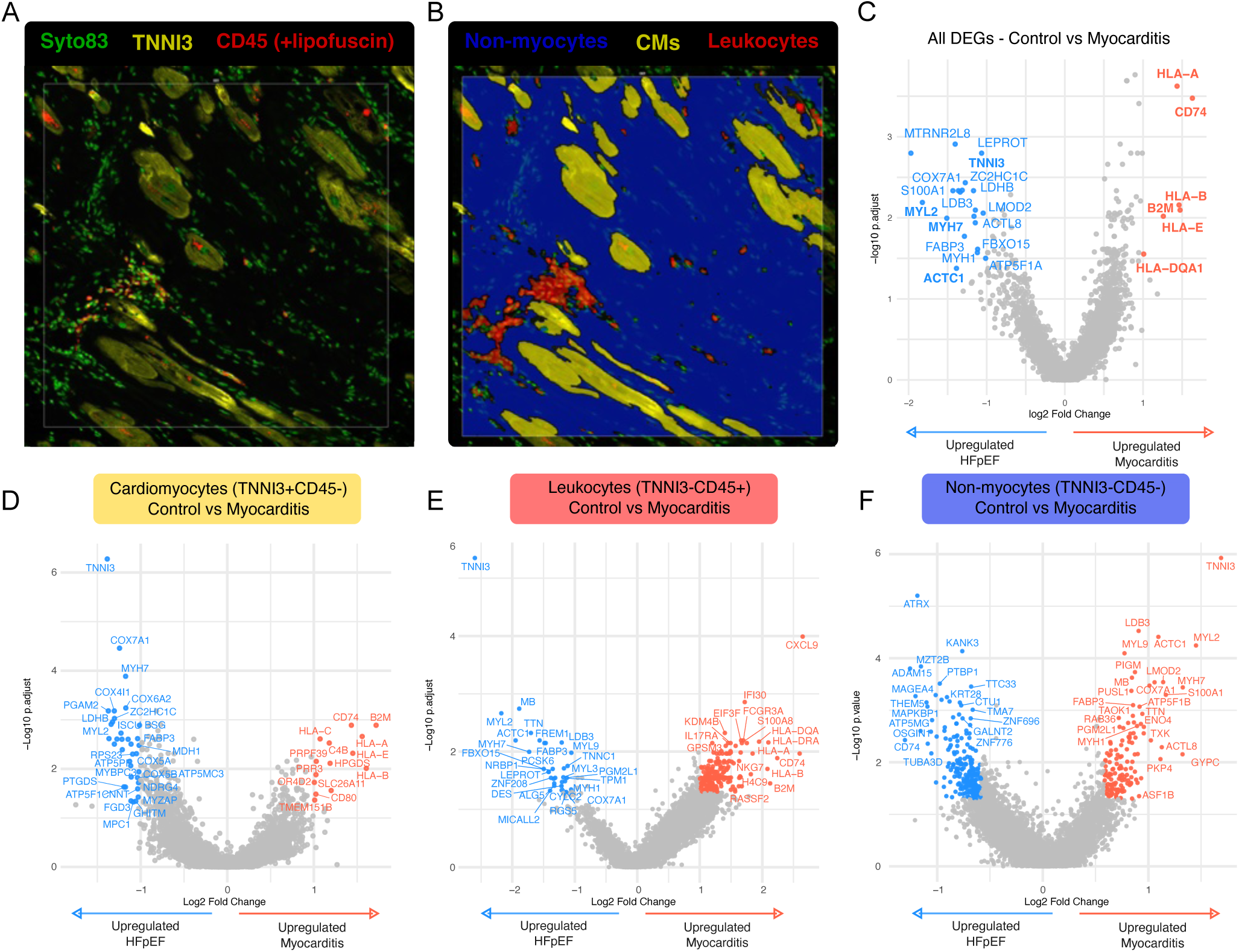
Compartment-specific spatial transcriptomic profiling of human myocarditis using GeoMx DSP supports prior regional signatures. **(A)** Representative immunohistochemical (IHC) micrograph of endomyocardial biopsy (EMBx) tissue highlighting cardiomyocytes (TNNI3⁺, yellow), leukocytes (CD45⁺, red), and nuclei (Syto83, green). **(B)** Representative segmentation overlay into three compartments: cardiomyocytes (TNNI3⁺CD45⁻, yellow), leukocytes (TNNI3⁻CD45⁺, red), and non-myocytes (TNNI3⁻CD45⁻, blue), for IHC-guided transcriptomics (GeoMx DSP). **(C)** Volcano plot showing all DEGs between controls and myocarditis in all segments, **(D)** TNNI3^+^CD45^-^ cardiomyocytes, **(E)** TNNI3^-^CD45^+^ leukocytes, and **(F)** TNNI3^-^ CD45^-^ non-myocytes/stromal cells. DEGs were computed using Q3 normalization followed by linear mixed-effects modeling with a FC threshold > 1.5 and an adjusted *p* < 0.05. For non-myocyte comparisons (F), unadjusted *p*-values were used due to lower segment counts and limited detection sensitivity.

To estimate cell-type composition within spatial transcriptomic segments, we applied SpatialDecon (v1.18.0), a reference-based deconvolution method designed for GeoMx data^10^. Using the human cell profile matrix ‘Heart_HCA.RData’^11^ cell type deconvolution was performed for each AOI to validate segmentation and confirm compartment-specific cellular identity. As expected, TNNI3⁺CD45⁻ segments were highly enriched for cardiomyocytes (Supplementary Figure 4A), while double-negative (TNNI3⁻CD45⁻) segments exhibited considerable heterogeneity, comprising diverse stromal populations such as fibroblasts, endothelial cells, and pericytes (Supplementary Figure 4B). Leukocyte-rich segments (TNNI3⁻CD45⁺) were primarily comprised of myeloid and lymphoid populations, consistent with spatial immunostaining and supporting effective immune compartment segmentation (Supplementary Figure 4C).

Differential gene expression was performed between control and myocarditis samples. First, all compartments were considered (Figure 4C, Supplementary Table 9). This illustrated that myocarditis is characterized by robust, widespread upregulation of antigen presentation and inflammatory genes, consistent with 10X Visium data. Notably, genes such as *HLA-A*, *B2M,* and *CD74* were upregulated in myocarditis relative to controls, aligning with Visium findings. Several transcripts were enriched in controls—including *MYH7*, *S100A1*, and *ACTC1.* These were also consistent with those identified in Visium datasets (Figure 4C, Supplementary Table 9).

We then performed differential gene expression analysis within each IHC-segmented compartment. Transcripts within cardiomyocytes (TNNI3⁺CD45⁻) from myocarditis samples demonstrated marked upregulation of antigen presentation genes including *HLA-A*, *HLA-B*, *HLA-E*, and *B2M*, alongside immune regulators *CD74* and *CD80* (Figure 4D, Supplementary Table 9). These transcripts encode core components of the MHC class I complex and associated antigen-processing machinery, implicating an active role for cardiomyocytes in immune signaling. Conversely, control samples showed increased expression of structural and metabolic genes, including *MYH7*, *COX7A1*, and *LDHB*. Interestingly, *TNNI3* was significantly augmented in controls relative to myocarditis, reflecting that HFpEF control patients may have increased cardiomyocyte specific *TNNI3* transcripts compared to myocarditis patients. These findings align with our 10X Visium data, reinforcing that cardiomyocytes in myocarditis shift toward an immunologically active state.

In myocardial leukocytes (TNNI3⁻CD45⁺), pronounced upregulation of *CXCL9, HLA-A* and *CD74* was observed in myocarditis samples relative to control segments (TNNI3^-^CD45^-^, Figure 4E, Supplementary Table 9), indicating heightened antigen presentation and chemokine-driven immune cell recruitment in immune foci and interstitial leukocytes. In contrast, controls exhibited minimal expression of these aforementioned genes, underscoring the distinct inflammatory profile observed in myocarditis compared to controls.

In non-myocyte segments (TNNI3⁻CD45⁻), we had comparatively low abundance of non-myocyte cells and reduced transcript detection. Statistical power was limited, necessitating the use of unadjusted p-values in differential expression analysis. Under these conditions, myocarditis samples exhibited upregulation of cardiomyocyte-structural genes including *TNNI3*, *MYH7*, *MYL2*, and *ACTC1*. This may reflect atypical expression of contractile machinery within non-myocyte populations, a finding previously reported in rodent studies^29^. However, in this population, we were unable to effectively exclude signal bleed-through from adjacent cardiomyocytes (with high transcript expression) from low signal regions, which may influence observed expression patterns (Figure 4F, Supplementary Table 9). We also observed elevated expression of *S100A1* and *LDB3*, suggesting aberrant calcium handling activity and cytoskeletal reorganization. Conversely, controls showed increased expression of nuclear and cytoskeletal regulatory genes, such as *ATRX*, *MZT2B*, *PTBP1*, and *ADAM15*, consistent with modulation of non-contractile cellular functions. Indeed, these genes are linked to cytoskeletal regulation and chromatin remodeling^30,31^—processes implicated in murine HFpEF models—though their specific upregulation in HFpEF remains to be confirmed.

### Cross-platform concordance and predicted ligand–receptor networks implicate cardiomyocyte antigen presentation in myocarditis

To identify the most robust findings from our work, we compared DEGs identified from myocarditis samples using both 10X Visium and GeoMx DSP platforms. First, we evaluated all compartments, yielding a total of 121 and 389 distinct DEGs detected by GeoMx and Visium, respectively, with 14 common genes differentially regulated in myocarditis across both platforms (Figure 5A, Supplementary Table 10). These included immune and antigen presentation–related transcripts such as *HLA-A*, *HLA-DQA1*, *IFI30* and *B2M*. Restricting the analysis to cardiomyocyte-enriched regions across both platforms modestly decreased concordance, revealing 192 and 137 distinct genes (GeoMx vs. Visium) and 10 shared DEGs between platforms (Figure 5B, Supplementary Table 10), including *HLA-A*, *HLA-E*, *B2M*, and *CD74* further reinforcing modulation of antigen presentation-related genes within cardiomyocytes in human myocarditis. To further investigate the potential role of these DEGs within immune communication networks, we constructed a ligand–receptor interaction map using OmniPath^17^ (Figure 5C). Ligand–receptor interactions were filtered to retain only those involving at least one gene from the union of overlapping DEGs identified in GeoMx and Visium analyses (Figure 5A–B).

**Figure 5.**
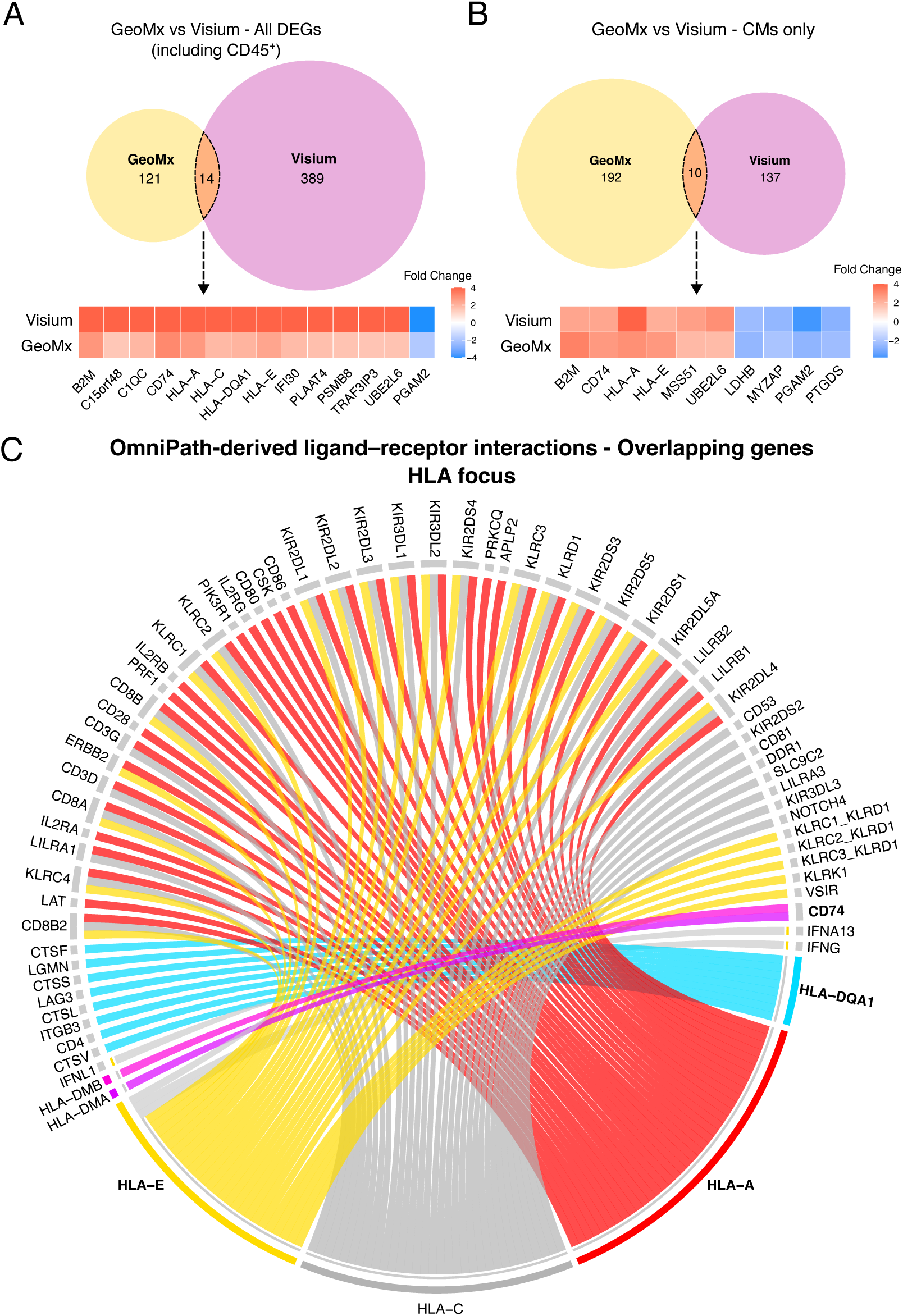
Cross-platform comparison of differential gene expression in myocarditis using GeoMx DSP and 10X Visium. **(A)** Proportional Venn diagram showing the overlap of DEGs between Visium (FindAllMarkers) and GeoMx datasets (all compartments). Fourteen DEGs were consistently differentially regulated in myocarditis relative to controls across both platforms. Corresponding fold changes for these overlapping genes are shown in the heatmap below. **(B)** Proportional Venn diagram comparing DEGs identified only in cardiomyocyte-stained segments (TNNI3⁺CD45⁻) and leukocyte depleted, cardiomyocyte-enriched genes (Visium), revealing ten shared DEGs between both datasets. Fold change values for these overlapping genes are shown in the heatmap. Color intensity in the heatmaps reflects the magnitude of absolute fold change values for each gene. Genes shown were filtered based on adjusted p-value of at least < 0.01 and exhibited consistent directionality of effect across platforms. Heatmap values for upregulated genes with FC higher than 4 were capped to this maximum value to aid visualization (see Supplementary Table 10 for values). **(C)** Chord plot illustrating inferred ligand– receptor interactions derived from differentially expressed genes in cardiomyocyte-enriched regions from both experimental techniques, focusing on overlapping antigen presentation–related genes, weighted by expression confidence. Arcs represent predicted interactions between ligands and immune receptors. Interactions were inferred using the OmniPath^17^ ligand–receptor database, and visualized using network-based filtering of curated, directional signaling interactions. Bolded genes represent overlapped genes present in OmniPath, between the two orthogonal experimental techniques.

Subsequently, chord diagram visualization highlighted *HLA-A*, *HLA-E*, and *HLA-DQA1* as major receptor hubs, interacting with immune cell ligands including *CD3D*, *CD8A*, *CD80*, and several *KIR* (NK-cell) family members (Figure 5C). Notably, *HLA-DQA1* exhibited a distinct interaction profile compared to *HLA-E* and *HLA-A*, by selective engagement with NK-associated receptors and molecules relevant to CD4⁺ T cell responses. Moreover, CD74, a class II MHC-associated chaperone, also emerged as a central signaling node for other HLA genes not in this dataset. Together, these findings indicate that cardiomyocytes in myocarditis undergo transcriptional induction of class I antigen presentation machinery and express receptor components capable of engaging in defined ligand– receptor interactions with infiltrating immune cells—which themselves upregulate class II molecules— implicating direct molecular crosstalk between parenchymal and immune compartments in the context of human myocarditis.

## DISCUSSION

This study presents the first comprehensive spatial transcriptomic analysis of endomyocardial biopsies in human myocarditis. We found that myocarditis is characterized by widespread transcriptional dysregulation beyond immune cell foci, providing compelling evidence that myocarditis is not a purely focal disease but a diffuse molecular state of global myocardial inflammation. Notably, both 10X Visium and GeoMx DSP platforms consistently demonstrated upregulation of antigen presentation-related genes in non-immune, cardiac cells—particularly cardiomyocytes. This suggests that cardiomyocytes actively participate in immune signaling in the context of human myocarditis.

Myocarditis has been classically considered a focal disease. Endomyocardial biopsy-based diagnosis of myocarditis relies on the identification of focal immune infiltrates^4–6^. Cardiac MRI based diagnosis is frequently used in the clinical setting, but diagnostic criteria developed for MRI are based on the understanding that the disease manifests in a patchy, multifocal pattern^32^. Our tissue transcriptomics analysis suggests that myocarditis is instead a global disease of the myocardium. This stands to reason, given that cardiomyocytes have been shown to secrete cytokines and respond to an array of immune signals^33–35^. Hence, this suggests that assessment of endomyocardial biopsies based on the expression of diffusely upregulated genes in myocarditis might achieve higher sensitivity than current histology-based assessment of endomyocardial biopsies, which has a high level of discordance—even among expert pathologists^36^.

Diagnosis of myocarditis is particularly challenging in clinical practice, in cases without overt myocardial injury, as these are classically labelled as borderline myocarditis^36^. We present the first spatial transcriptomics analysis of borderline myocarditis. Analyzing borderline myocarditis, we found that this diagnostic category likely encompasses: 1) misclassified cases from lack of compelling features of myocarditis, (2) cases that are inappropriately interpreted as a pathological deviation from healthy hearts and (3) cases that present a lower level of inflammatory activation than definitive cases of myocarditis. Presently, our data cannot determine whether the observed reduction in immune activation is indicative of early disease captured along a continuum from control to myocarditis, or if it represents an entirely different pathological state. However, we noted that myocarditis was associated with clear upregulation of antigen presentation-related genes when compared to borderline myocarditis. This suggests that antigen presentation by cardiomyocytes might be a determining factor in the progression from borderline myocarditis to myocarditis.

The most robust and reproducible finding of our spatial transcriptomic analyses of human endomyocardial biopsies was the upregulation of MHC class I and II-related molecules in myocarditis across both leukocyte-rich and leukocyte-depleted compartments. Comparison of Visium and GeoMx datasets revealed a distinct subset of overlapping DEGs, enriched for immune and antigen presentation genes, which were consistently expressed in cardiomyocytes from myocarditis patients. These included *HLA-A*, *HLA-DQA1*, *B2M*, *CD74*, and *IFI30*. Consistently, ligand–receptor interaction modeling using OmniPath, filtered to genes overlapping between GeoMx and Visium datasets, identified *HLA-A*, *HLA-E*, and *HLA-DQA1* as prominent receptor nodes, each with multiple predicted interactions involving immune ligands. Cytokines such as IFN-γ are known to induce MHC class I expression in cardiac tissue^37,38^. Several IFN-γ related genes were also upregulated in this dataset, particularly *IFI30* and *B2M*, which support the functional antigen-presenting capacity of cardiomyocyte-expressed HLA-A. Indeed, infection of murine ventricular myocytes with Coxsackievirus B3 induces upregulation of MHC-I and, to some extent, also of MHC-II molecules^34^. Moreover, a recent tissue transcriptomics study of murine reoviral myocarditis reported upregulation of genes involved in antigen presentation (in the absence of MHC transcripts) in endothelial cells but not in cardiomyocytes^39^. Nearly three decades ago, an immunohistochemical study of human endomyocardial biopsies reported expression of MHC-I and MHC-II molecules on cardiomyocytes in the context of myocarditis^40^. Our findings reinforce this observation, supporting the notion that human myocarditis is characterized by upregulation of MHC-I and possibly MHC-II molecules on cardiomyocytes and adjacent cells, expanding on the emerging notion that non-immune cardiac cells are capable of presenting antigens^41^.

Upregulation of HLA-A in cardiomyocytes in myocarditis may facilitate recognition by CD8⁺ T cells, potentially leading to immune-mediated cardiac injury. With respect to the HLA-DQ complex (classical MHC class II molecule), α-chain *HLA-DQA1* heterodimerizes with the β-chain *HLA-DQB1*, allowing for antigen presentation to CD4⁺ T cells^42^, suggesting a likely role for cardiomyocytes in initiating or sustaining helper T cell–mediated immunity. In addition, CD4⁺ T cells specific for cardiac antigens have been shown to contribute to disease progression in both pressure overload–induced heart failure and autoimmune myocarditis^43,44^. Notably, CD74, a key MHC class II chaperone, also emerged as a prominent signaling node, further supporting the engagement of functional antigen processing pathways in cardiomyocytes in myocarditis. Although expressed at low levels in cardiomyocytes under baseline conditions, CD74 contributes to LPS-induced cardiac dysfunction *in vivo* by promoting inflammatory signaling pathways that disrupt calcium homeostasis and impair contractile function^45^. Further investigations are warranted to determine whether cardiomyocyte-mediated antigen presentation represents a viable therapeutic target to treat myocarditis, and whether class II is directly expressed by cardiomyocytes.

Several limitations of our study merit consideration. First, although this represents the largest spatial transcriptomic study of human myocarditis to date, we included 38 endomyocardial biopsies, leaving the overall cohort size modest, which may limit statistical power for detecting subtle transcriptional differences, particularly in rare, undetectable cell populations. Second, we intentionally included a heterogeneous group of patients to investigate molecular pathways common to myocarditis, beyond subtypes of the disease. While this served our purpose and helped us distil the molecular pathways most relevant to myocarditis, the heterogeneity of our cohort in the context of our limited cohort size is a potential source of systemic variation. Third, despite the implementation of rigorous pre-processing quality control and cross-platform validation, reliance on archival FFPE tissue introduces inherent variability in RNA integrity and transcript capture efficiency. Fourth, spatial resolution is constrained by the spot or area-of-interest size inherent to Visium and GeoMx platforms, limiting our ability to achieve true single-cell resolution and fully resolve cellular heterogeneity within densely populated regions. While cardiomyocyte enrichment and IHC-guided segmentation improved compartmental resolution, cell-type deconvolution remains approximate. Fifth, while transcriptomic evidence suggests antigen presentation by non-immune cells, and our findings are in line with historical findings obtained via IHC on human biopsies^40^, direct protein-level validation (e.g., IHC or spatial proteomics) was not performed in this study. Lastly, the limited concordance between spatial platforms is noteworthy, which likely reflects differences in gene panel design, statistical power, and spatial sampling strategy — with GeoMx relying on specific ROIs and Visium capturing entire endomyocardial biopsy sections.

## Conclusion

Taken together, this study represents the first and most substantial tissue transcriptomic analysis of human myocarditis and borderline myocarditis to date, establishing a novel molecular framework for understanding myocarditis as a spatially diffuse and transcriptionally dynamic process, rather than confined to discrete histological lesions. By integrating dual-platform spatial transcriptomics with compartment-specific analysis, we show that in human myocarditis antigen presentation and immune activation extend into non-immune cardiac cells, challenging classical paradigms of immune dynamics in myocardial disease. Beyond redefining myocarditis tissue pathology, these findings underscore the potential of spatial transcriptomics to inform next-generation diagnostic and therapeutic approaches. Future work should extend these insights through higher-resolution spatial profiling, longitudinal sampling, and functional validation to clarify the diagnostic value, causal roles, and potential therapeutic significance of these molecular programs in disease progression and recovery.

## Supporting information

Cohen_etal_2025_Clinical_Myocarditis_Supplemental tables

## ACKNOWLEDGEMENTS

This study was partially funded by a grant from the W.W. Smith Charitable Trust and by a Catalyst Grant awarded to Dr. L. Adamo by the Johns Hopkins Division of Cardiology. Dr. C.D. Cohen was supported by American Heart Association Postdoctoral Research Fellowship (Grant #: 25POST1361888, awarded 2025). We also acknowledge Russell Hughes from the Single Cell & Transcriptomics Core, Institute for Basic Biomedical Sciences, Johns Hopkins University for providing technical and experimental expertise to this project.

## FIGURE LEGENDS – SUPPLEMENTAL MATERIAL

**Supplementary Figure 1.**
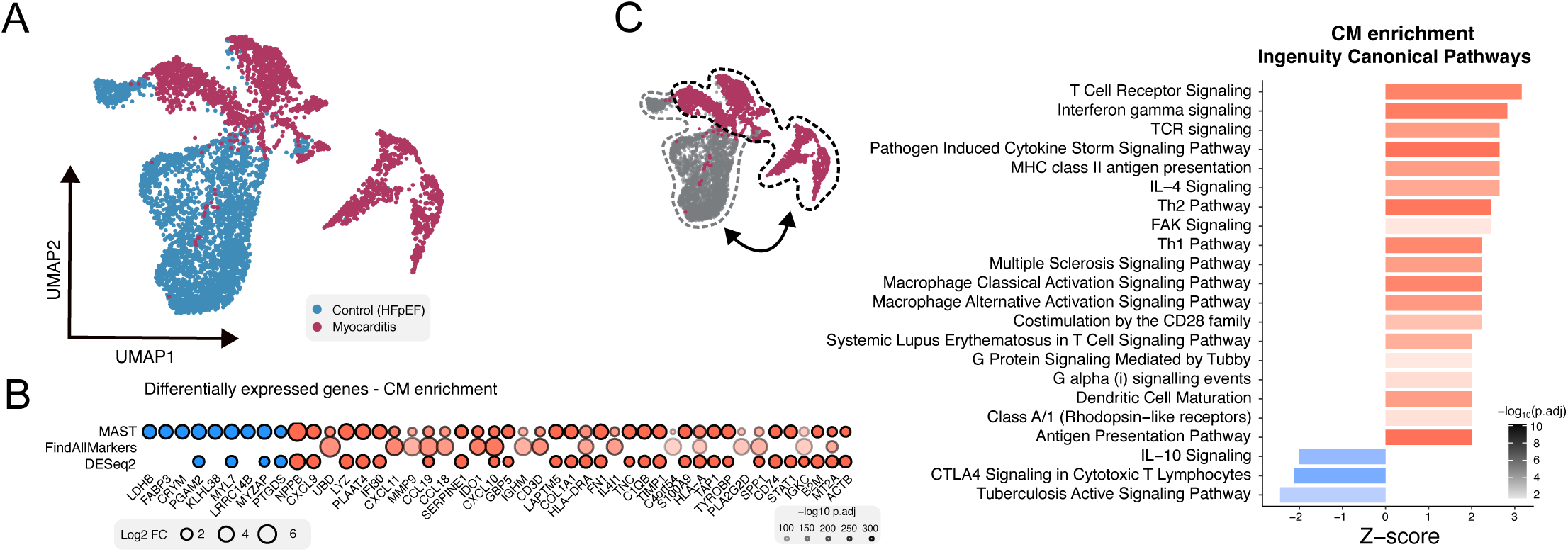
Cardiomyocyte-enriched transcriptional changes in myocarditis. **(A)** UMAP projection of cardiomyocyte-enriched spatial transcriptomic spots from myocarditis (red) and control (HFpEF, blue) samples, showing robust separation between disease states. **(B)** Dot plot showing the top differentially expressed genes between myocarditis and controls across three statistical methods: MASTcpmDETrate, FindAllMarkers, and DESeq2. Dot size represents the absolute log2 fold change, and color indicates adjusted p-value significance (−log10 scale). **(C)** Top canonical pathways identified from Ingenuity Pathway Analysis (IPA) of cardiomyocyte-enriched spots using FindAllMarkers-derived differentially expressed genes. Pathways are ranked by z-score, with red indicating upregulated and blue indicating downregulated pathways in myocarditis compared to controls. Highlighted pathways reflect strong immune activation and antigen presentation programs in cardiomyocytes.

**Supplementary Figure 2:**
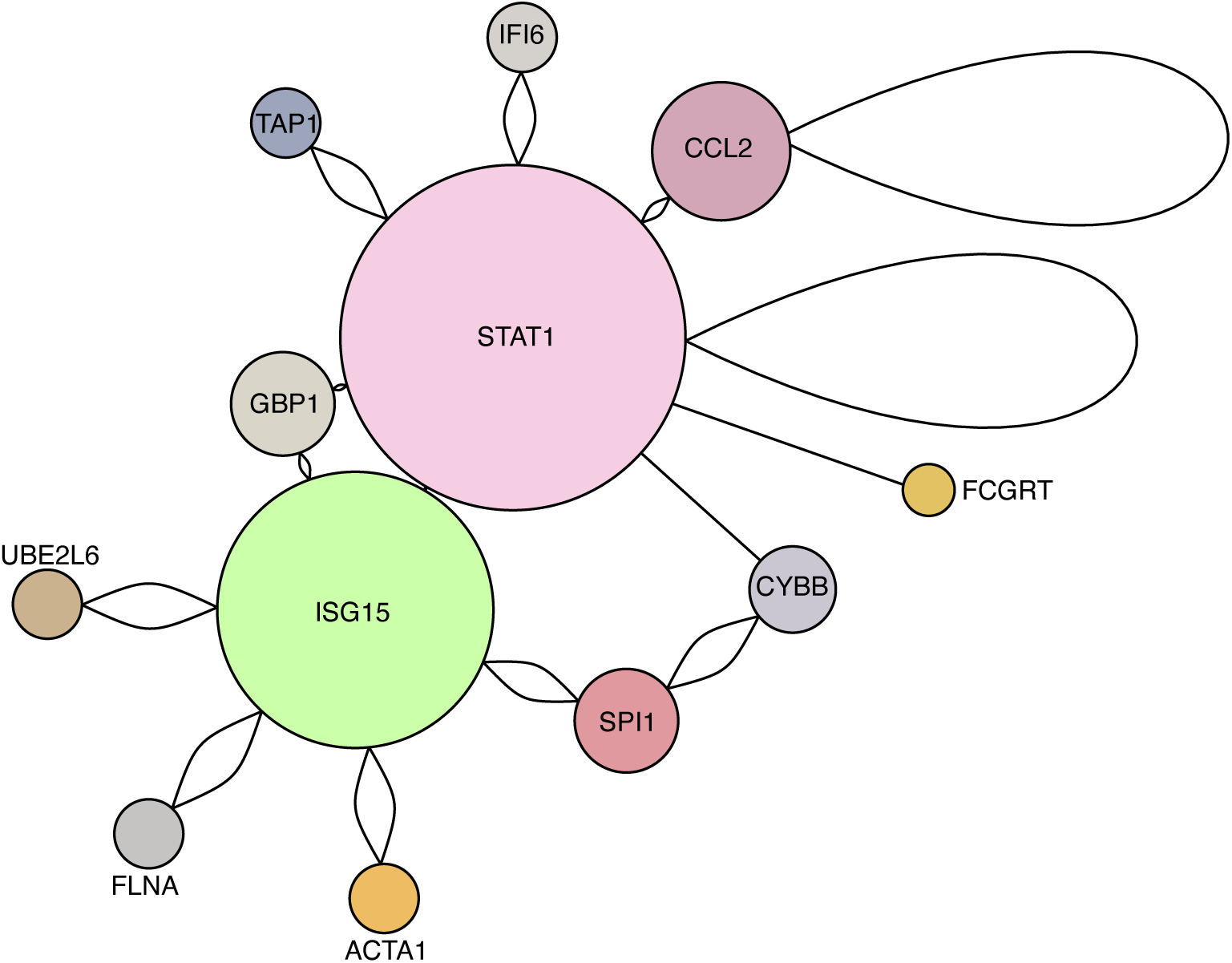
EMBx samples refined for CMs show central role for STAT1 signaling in myocarditis. Protein–protein interaction (PPI) network generated from DEGs identified in the Visium spatial transcriptomics dataset using the FindAllMarkers output. Interactions were curated through the InnateDB database, highlighting innate immune signaling relationships among genes enriched in myocarditis samples. Node size reflects degree centrality, with larger nodes representing more connected hub genes. STAT1 and ISG15 emerged as central network hubs, interacting with multiple immune-associated genes including *CCL2*, *IFI6*, *GBP1*, and *CYBB*. Edges denote experimentally validated or high-confidence curated interactions.

**Supplementary Figure 3:**
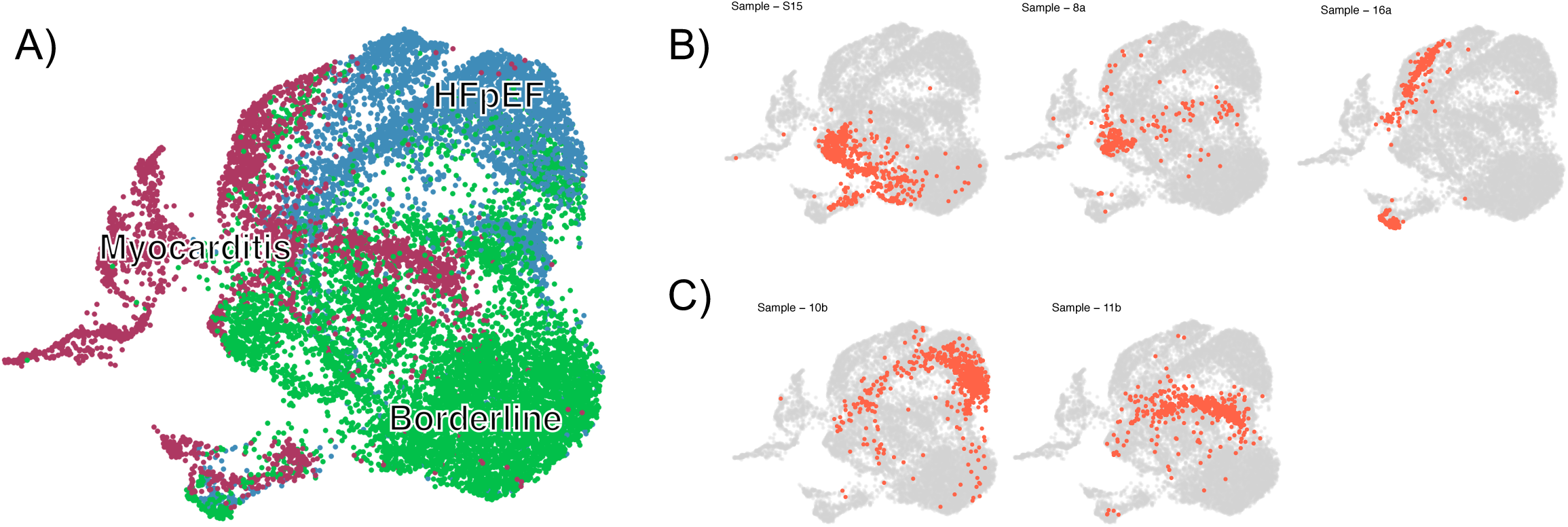
Reassignment of borderline myocarditis samples based on transcriptional overlap with HFpEF and myocarditis clusters. **(A)** UMAP projection illustrating borderline myocarditis spots (green) clustering with either HFpEF (blue) or myocarditis (pink), suggesting transcriptional misalignment with the borderline classification. **(B–C)** Individual borderline samples highlighted on the UMAP, showing overlap with either myocarditis or HFpEF transcriptomic profiles. These samples were excluded from downstream analyses to reduce diagnostic noise.

**Supplemental Figure 4.**
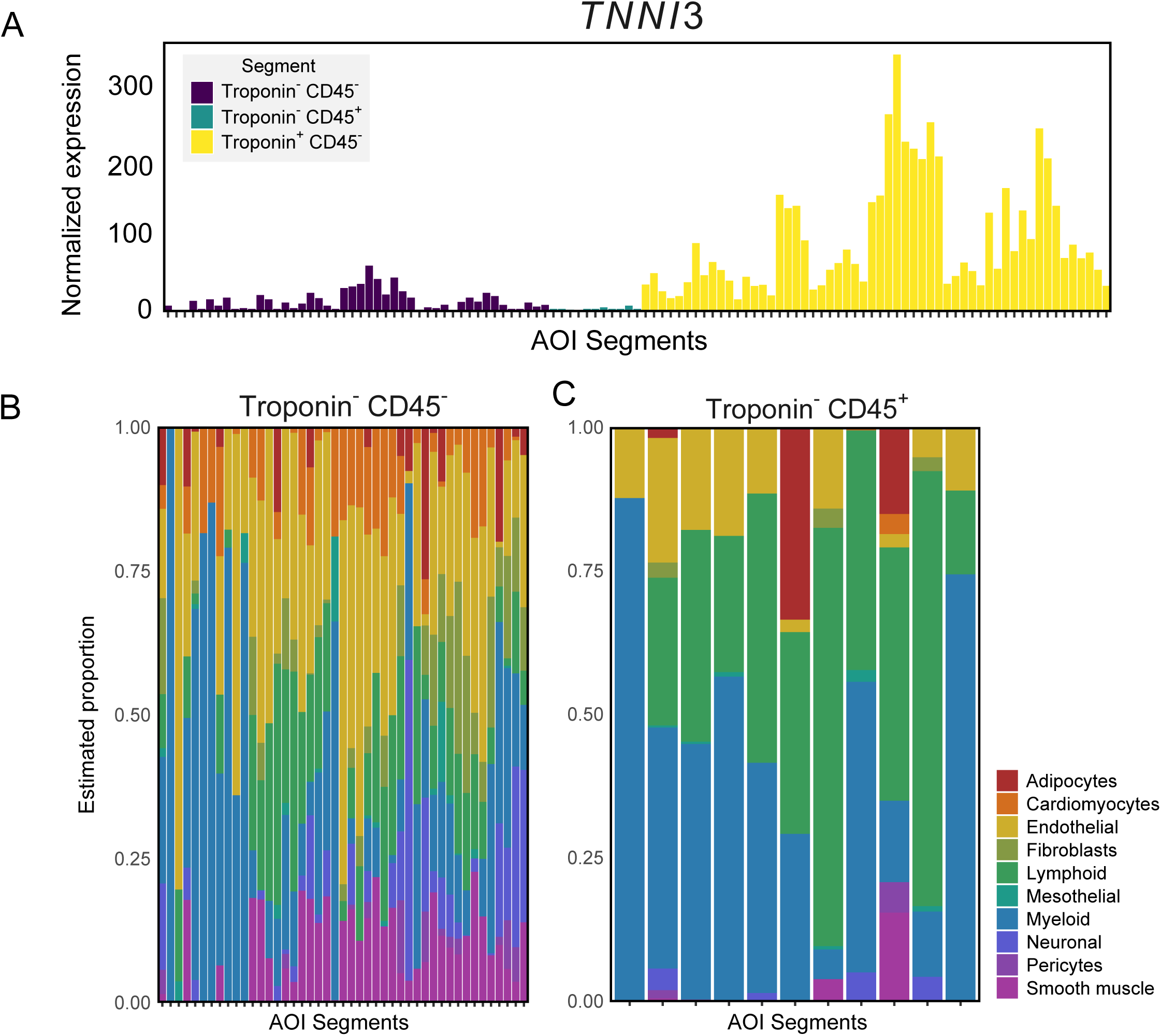
Validation of compartment-specific segmentation in GeoMx DSP data. (A) Expression of TNNI3 (Troponin I type 3) across segmented Areas of Interest (AOIs), showing highest expression in TNNI3⁺CD45⁻ segments (yellow), consistent with cardiomyocyte enrichment. (B–C) Cell-type deconvolution analysis of TNNI3⁻CD45⁻ (non-myocyte) and TNNI3⁻CD45⁺ (leukocyte) compartments reveals expected enrichment of fibroblasts, endothelial cells, lymphoid and myeloid populations, validating IHC-based spatial compartmentalization. TNNI3^-^CD45^+^ AOI segments only collected from myocarditis sections due to the lack of immune infiltrate in control samples. These data support the specificity of segmentation strategy used for downstream transcriptomic analyses.

