## Supplementary material for "Tissue transcriptomics of endomyocardial biopsies reveals widespread molecular perturbations independent of leukocyte-rich foci in human myocarditis": Cohen_etal_2025_Clinical_Myocarditis_Supplemental tables: Table 1. Clinical Characteristics of Patients Included in the Study_v4.docx

### Table 1. Summary Statistics by Diagnostic Group

| Diagnosis | Sex - Female (Total) | Median age  (± SD) | Median LVEF (± SD) | Median NT-proBNP (± SD) |
| --- | --- | --- | --- | --- |
| Control | 6 (12) | 67.5 ± 13.2 | 65.0 ± 11.1 | 258.5 ± 894.0 |
| Myocarditis | 7 (10) | 46.5 ± 24.8 | 37.5 ± 22.3** | 7931.0 ± 7712.0** |
| Borderline | 3 (12) | 59.0 ± 21.9 | 60.0 ± 4.2 | 3707.0 ± 4135.0 |

Summary of patient characteristics at time of endomyocardial biopsy, grouped by final diagnostic classification: control, myocarditis, and borderline myocarditis. Sex is shown as number of female patients over total patients per group. The other values represent medians with standard deviations (± SD). LVEF values were derived from ranges indicated in clinical reports using the midpoint if an exact value was not available. NT-pro-BNP is expressed in pg/ml and LVEF is expressed in %. Both variables showed statistically significant differences between HFpEF and myocarditis. Age was not significantly different across groups. * *P* < 0.05, ***P* < 0.01 – HFpEF vs. Myocarditis. †*P* < 0.05 HFpEF vs borderline. Statistical significance was determined by one-way ANOVA and Kruskal–Wallis *post-hoc* test.
